# Molecular insights into peptide agonist engagement with the PTH1 receptor

**DOI:** 10.1101/2022.09.04.506565

**Authors:** Brian P. Cary, Elliot J. Gerrard, Matthew J. Belousoff, Madeleine M. Fletcher, Yan Jiang, Isabella C. Russell, Sarah J. Piper, Denise Wootten, Patrick M. Sexton

## Abstract

The parathyroid hormone (PTH) 1 receptor (PTH1R) is a class B1 G protein-coupled receptor (GPCR) that critically regulates skeletal development and calcium homeostasis. Despite extensive study, the molecular underpinnings of PTH1R stimulation by its cognate hormones, as well as by therapeutic agents, remain unclear. Here, we describe cryo-EM structures of the PTH1R in complex with active fragments of the two hormones, PTH and parathyroid hormone related protein (PTHrP), the peptidic drug abaloparatide, as well as the engineered tool compounds, long-acting PTH (LA-PTH) and the truncated peptide, M-PTH(1-14). We found that the N-terminus of each agonist that is critical for activity, engages the transmembrane bundle in a topologically similar fashion, which reflects similarities in measures of Gαs activation. The full-length peptides bind the extracellular domain (ECD) using a shared interface but induce subtly different ECD orientations relative to the transmembrane domain (TMD). In the structure bound to M-PTH, an agonist which only binds the TMD, the ECD is completely unresolved, demonstrating that the ECD is highly dynamic when unconstrained by a peptide. High resolutions enabled identification of water molecules near the peptide and G protein binding sites, some of which are structurally conserved with other class B1 GPCRs. Our results shed light on the action of orthosteric agonists of the PTH1R and provide a foundation for structure based-drug design.

## INTRODUCTION

GPCRs are critical conduits for signal transduction and comprise the single largest family of proteins within the human genome. Primarily expressed in renal and bone tissue, the PTH1R, a class B1 GPCR, maintains circulating calcium and phosphorus homeostasis (Martin, Sims and Seeman, 2021). The PTH1R is activated by two polypeptides hormones, parathyroid hormone (PTH) and parathyroid hormone-related protein (PTHrP). The full-length, mature hormones are 84 and 141 amino acids in length, respectively, but the N-terminal fragments (∼34 amino acids) are sufficient to elicit activity (Cheloha *et al*., 2015). Genetic studies in rodents showed divergent effects for knockout of PTHrP or PTH, revealing greater relevance for pre- and post-natal physiology, respectively (Martin, Sims and Seeman, 2021). While both hormones are similarly potent agonists for eliciting cAMP production downstream of Gαs activation, the primary PTH1R signaling pathway, PTH and PTHrP display distinctions in their preference for binding the G protein-uncoupled receptor state (Hattersley *et al*., 2016) and in duration of cAMP signaling (Ferrandon *et al*., 2009).

Due to its physiological importance, impairments in PTH1R signaling are involved in several human diseases (Cheloha *et al*., 2015, p. 1). Tumor-derived PTHrP production promotes adipose and muscle tissue wasting (Kir *et al*., 2014). Hypoparathyroidism and autoantibodies blocking PTH activity can result in serum hypocalcemia (Mandl *et al*., 2021). Conversely, constitutive activity inducing mutations in PTH1R can lead to hypercalcemia and profound skeletal dysplasia (Schipani *et al*., 1996). Beyond its capacity to contribute to disease, the PTH1R is a validated target for clinical interventions. Full-length, recombinant PTH is used for the treatment of hypoparathyroidism (Kim and Keating, 2015), and the N-terminal 1-34 fragment of PTH is the active ingredient in the osteoporosis drug, teriparatide. Moreover, abaloparatide (ABL), an analogue of PTHrP that shows superior bone building activity compared to teriparatide (Leder *et al*., 2015), is approved for the treatment of osteoporosis.

The PTH1R, like all class B1 GPCRs, contains a 7-helix transmembrane domain (TMD) and a relatively large extracellular domain (ECD), both of which are required for high affinity binding of endogenous peptides. Alone, the PTH1R ECD is soluble and retains the ability to bind the C-terminal region of peptide agonists. Three crystal structures of the isolated PTH1R ECD have been reported (Pioszak and Xu, 2008; Pioszak *et al*., 2009, 2010): one bound to PTH, one bound to PTHrP, and an apo structure. These ECD crystal structures revealed that PTH and PTHrP adopt an extended helical formation to bind a shared hydrophobic grove of the ECD. Until recently, the structure of the peptide N-terminal region in its receptor bound form was unknown, but two co-structures with the full length PTH1R revealed that the agonist helical conformation continues from the C-terminus into the TM pocket (Ehrenmann *et al*., 2018; Zhao *et al*., 2019), and these TMD interactions are required to elicit transducer engagement and signal transduction (**Figure 1a**). However, in those structures, the agonists of PTH1R were engineered to support structural stability, and do not provide direct insights into the binding of the native hormones or therapeutics. To address this gap, we present single-particle cryo-EM structures of five peptide agonists bound the PTH1R and the Gs heterotrimeric G protein.

**Figure 1:**
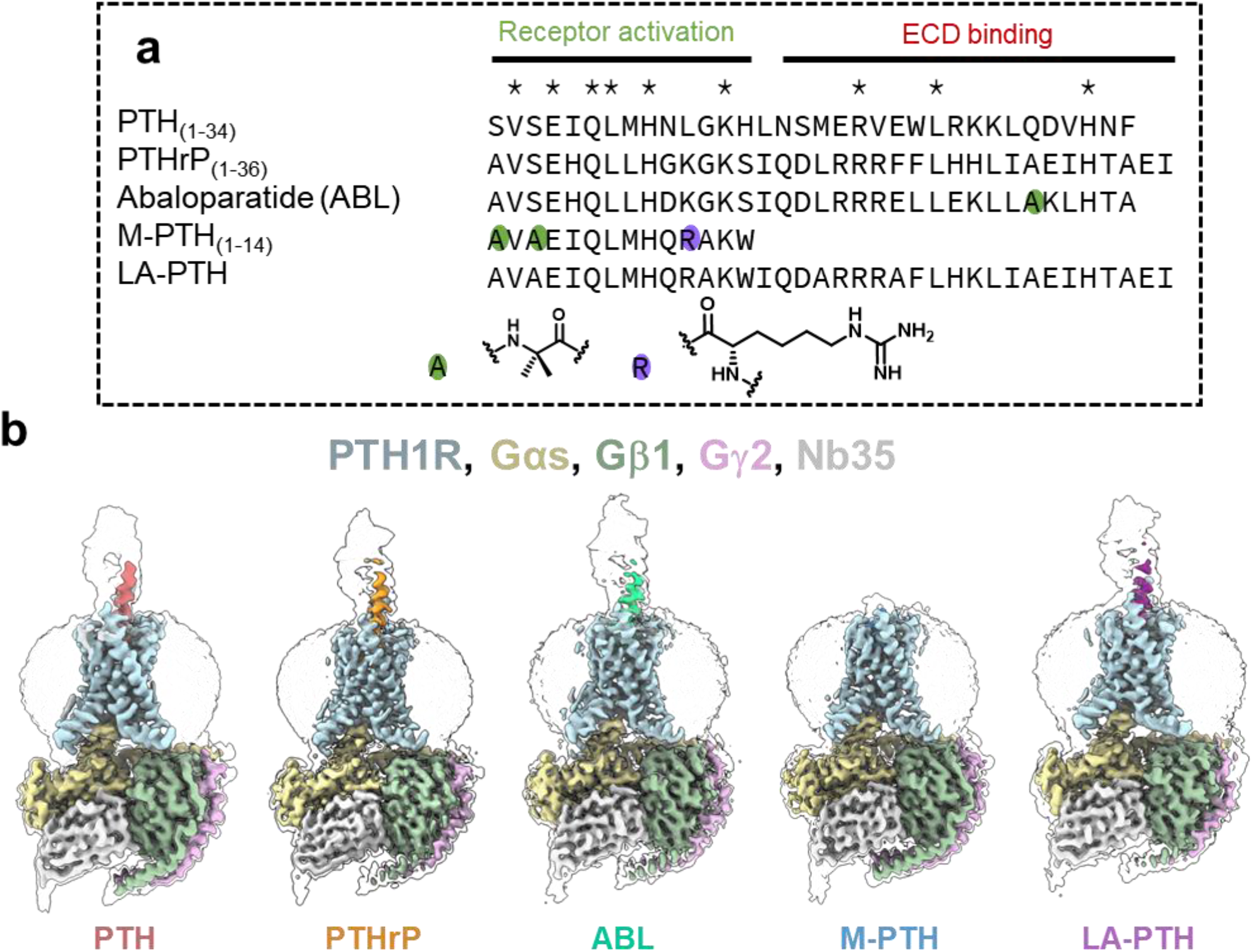
(**a**) Sequences of peptide PTH1R agonists. Asterisks indicate residues shared by the five peptides (or the 4 extended peptides for residues that are beyond the C-terminus of M-PTH). 2-amino butyric acid (AIB) residues are indicated with a green oval and homoarginine is indicated with a purple oval; the structures of these residues are shown. (**b**) Cryo-EM maps for the PTH1R bound to individual peptides, in complex with DNGαs, Gα2;1, Gα32 and Nb35 with map segments coloured according to the figure labelling. The colored surfaces show the map at high threshold, and the transparent silhouettes show the map at low threshold, allowing for visualization of the micelles and more dynamic ECDs.

## RESULTS

### Cryo-EM structures of peptides bound to PTH1R/Gαs

To gain a better understanding of the molecular mechanisms of PTH1R activation, we performed structural and conformational variance studies using single particle cryo-EM to determine structure and dynamics of the PTH1R bound to its canonical G protein (Zhao *et al*., 2019), Gs, in complex with five PTH analogues (**Figure 1a**). This included complexes with the drug abaloparatide (ABL) and the active fragments of the two native hormones, PTH(1-34) and PTHrP(1-36), due to their physiological and therapeutic importance, as well as the truncated peptide M-PTH(1-14) that selectively binds the PTH1R transmembrane domain, and the long-acting PTH analogue (LA-PTH) that can induce prolonged signaling (Noda *et al*., 2020; White *et al*., 2021). For conciseness, these peptides are referred to as PTH, PTHrP, M-PTH, and LA-PTH throughout the text and figures.

A construct encoding human PTH1R (**Figure S1**) was co-expressed with dominant negative Gαs (Liang *et al*., 2018), Gβ1 and Gγ2 in *Trichoplusia ni* insect cells. Following complex formation with nanobody35 (Rasmussen *et al*., 2011), saturating peptide, and apyrase treatment, PTH1R was solubilized in lauryl maltose neopentyl glycol/cholesteryl hemmisuccinate mixed micelles. Anti-FLAG affinity enrichment and size exclusion chromatography (SEC) were used to purify the complexes prior to vitrification and cryo-EM imaging (**Figure S2**). For each agonist, a substantial quantity of PTH1R dissociated from the G protein heterotrimer was present following affinity purification, which we were unable to eliminate from final samples by SEC (**Figures S2-S4**). However, image classification enabled isolation and reconstruction of the active-state receptor-G protein coupled complexes, highlighting the ability of single particle cryo-EM to contend with heterogenous samples.

Following image classification and *ab initio* reconstruction, we refined maps of PTH1R bound to PTH(1-34), PTHrP(1-36), ABL, M-PTH, and LA-PTH to gold-standard Fourier shell correlation (FSC, 0.143) resolutions of 2.55 Å, 2.94 Å, 3.09 Å, 3.03 Å, and 2.76 Å, respectively (**Figures 1b; S6**). Sufficient resolution was observed throughout the receptor and G protein to enable atomic modelling for most components. As observed for several class B1 GPCR structures, poor resolution for intracellular loop 3 (ICL3) and the G protein α-helical domain (AHD) was observed, which did not allow us to model these regions. There was also very limited density for ECL1, as has been previously reported for cryo-EM and crystal structures of the PTH1R (Ehrenmann *et al*., 2018; Zhao *et al*., 2019), consistent with high mobility in this region. Consensus backbone conformations for the receptor TMD, G protein, and peptide N-terminus were comparable among the structures containing different peptide agonists (**Figure 2d**), along with many of the interactions between peptide side chains and the receptor, and across the receptor G protein interface (**Table S1**) reflecting the pharmacological similarities of the peptides for PTH1R activation of Gαs; measured at the G protein level (Olsen *et al*., 2020) and production of secondary messenger (cAMP) (**Figures 2e-2f; S7**) (Shimizu, Guo and Gardella, 2001; Sato *et al*., 2021).

**Figure 2:**
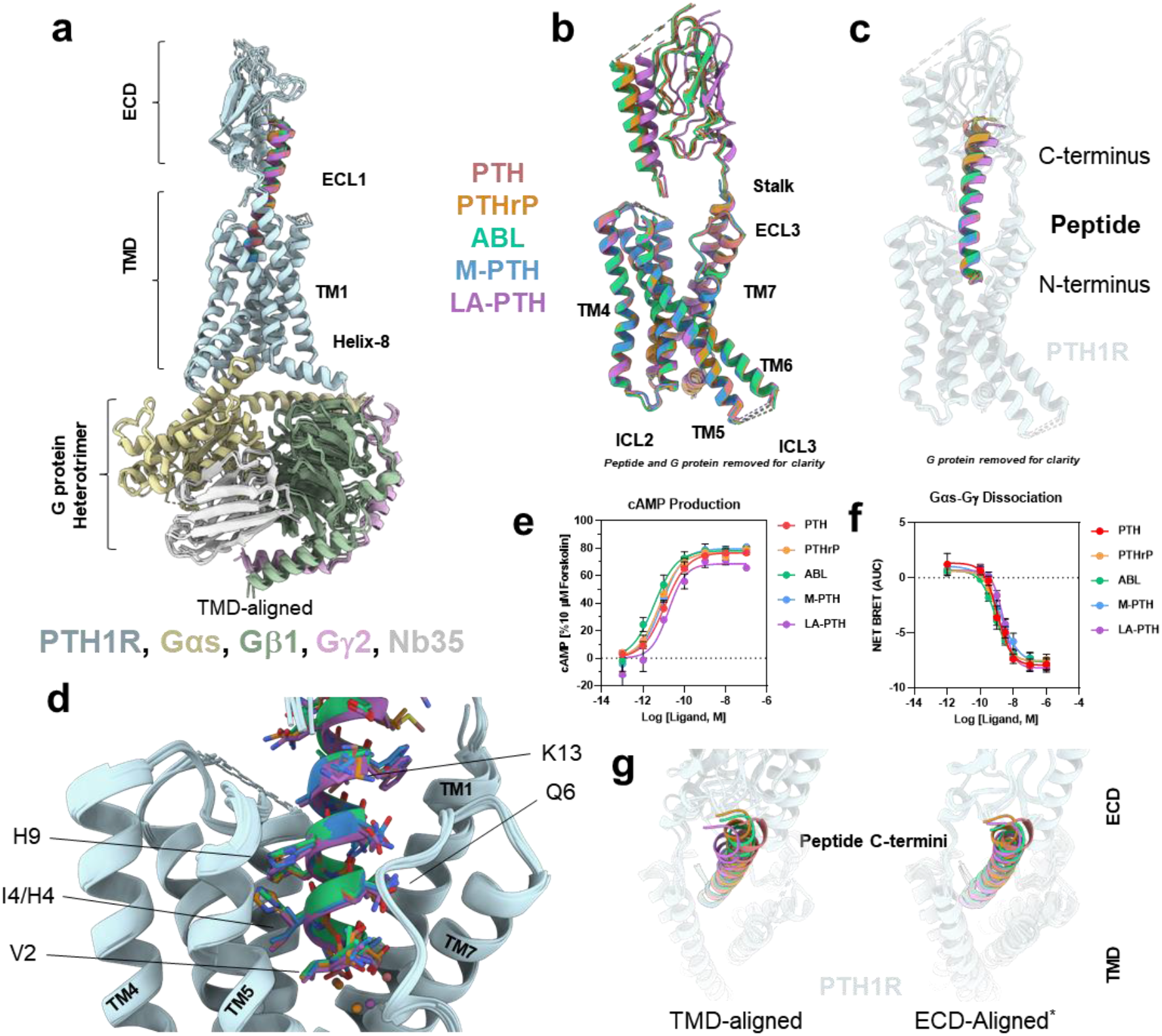
(**a**) The structures of PTH1R bound to Gs protein heterotrimer and diverse peptides. Models were aligned on the receptor TM domains (TMDs). Chains were colored according to the displayed keys. (**b**) Backbone structures of PTH1R with G proteins and peptides removed for clarity, aligned on TMDs. Models were colored according to the peptide to which each is bound. (**c**) Backbone structure of peptide agonists bound to the PTH1R colored according to the key. (**d**) Structures of peptides focused on the TMD pocket, aligned by the receptor TMD. (**e**) Peptide concentration-response for 3’,5’-cyclic adenosine monophosphate (cAMP) production in HEK293 cells transiently expressing the PTH1R measured as area under the curve (AUC) until 14 min after ligand addition. 100% represents the response in the presence of 10 µM forskolin. Data are mean ± SEM, N = 3. (**f**) Peptide concentration-response for dissociation of GαsL and Gα31 detected by bioluminescence resonance energy (BRET) in transiently transfected HEK293 cells, measured as AUC until 33 min after ligand addition. Data are mean ± SEM, N = 4. (**g**) A top view of the peptide C-termini aligned by receptor TMD (left) and extracellular domain (ECD) (right). ^*^excluding PTH1R bound to M-PTH, because the ECD is too poorly resolved to be modelled.

Despite sharing a common binding interface on the ECD, each of the full-length peptides induced subtly distinct orientations of the ECD with respect to the TMD (**Figure 2g**). The structures containing PTHrP and ABL were most comparable in ECD configuration, likely owing to their close sequence homology (**Figure 1a**). LA-PTH induced the most distinctive ECD arrangement with a ∼5 Å deflection of its C-terminus (measured by His32 Cα) towards TM1 compared to PTH. Correspondingly, the helix of LA-PTH is more bent than that of PTH. 3D-variability analysis (Punjani and Fleet, 2021) indicated a substantial degree of ECD and peptide motion for each of the full-length agonist-bound structures (**Video S1**), which agrees with the low local map resolution in this region (**Figure S6**). The motions observed using this analysis indicate that the peptide C-termini sample large and overlapping conformational space, but the consensus refinement indicates that the primary conformational preferences of the peptides are distinct.

In their critical N-terminal regions, six residues (∼40%) are conserved among the set of peptides we examined (**Figure 1a**). All of these analogous residues, with the exception of Lys13, are important for agonist activity in the context of PTH(1-14) (Luck, Carter and Gardella, 1999), and all bind similarly within the TMD (**Figure 3a; Table S1**). The sidechain of Val2 occupies a hydrophobic pocket formed by L292^3.40^, I367^5.43^, and L368^5.44^ (superscript numbers refer to the conserved class B1 GPCR numbering system of Wootten et al, (Wootten *et al*., 2013)). Conserved with several peptide hormones in the secretin family, Glu4 forms electrostatic interactions with R233^2.53^. Gln6 contacts the sidechain of Y429^ECL3^ and forms a hydrogen bond with Gln440^7.38^. Leu7 makes extensive hydrophobic interactions with the receptor mediated by F184^1.36^, L187^1.39^, Y191^1.43^, M441^7.39^, and M445^7.43^.

**Figure 3:**
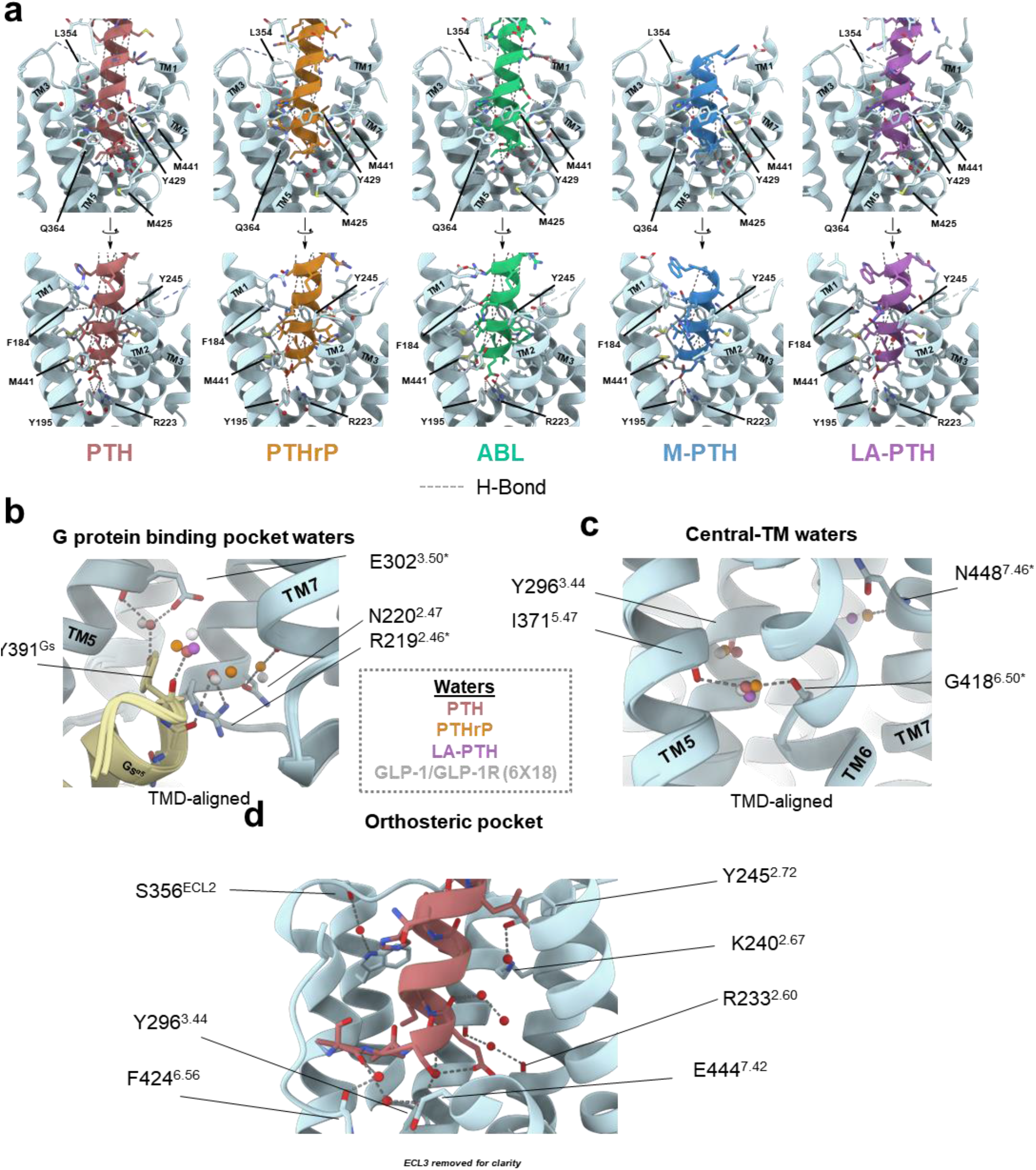
(**a**) Close-up side views of the PTH1R TMD bound to PTH, PTHrP, ABL, M-PTH, and LA-PTH (left to right). (**b-d**) Structural waters are shown as colored spheres (according to the bound peptide or structure in the key) in the vicinity of the G protein binding site (**b**), central transmembrane region (**c**), and PTH binding site (**d**). Select residues are displayed/labeled. Hydrogen bonds are show as dotted lines.

The identity of the residue at position 5 contributes to the different phenotypes exhibited by PTH and PTHrP; PTH shows a higher affinity for the G protein-uncoupled state and increased signaling duration compared to PTHrP (Dean *et al*., 2008; Yu *et al*., 2022). PTH, LA-PTH, and M-PTH contain an Ile residue at position 5, whereas PTHrP and ABL contain a His residue. A His5 → Ile5 substitution in PTHrP increased binding affinity and signaling half-life to levels comparable to PTH (Dean *et al*., 2008). The sidechain at position 5 inserts into a hydrophobic pocket that is enclosed by F288^3.36^, L289^3.37^, L292^3.40^, and I363^5.43^ (**Table S1**). This pocket may be more accommodating of a hydrophobic isoleucine sidechain than that of a histidine residue. In our map of the complex with PTHrP, we observed a weak density in this hydrophobic pocket near the δ carbon atom of PTH when the two models are aligned (**Figure S8**). This unmodeled density might correspond to a water that is unfavorably restrained in a hydrophobic pocket, potentially contributing to the lower affinity of PTHrP compared to PTH.

### The M-PTH-bound PTH1R ECD is highly mobile

M-PTH is unique among the agonists we examined because it is shorter in length and contains three non-native residues: one homoarginine and two 2-aminoisobutyric acid (Aib) residues. Due to the truncated sequence, M-PTH is thought to exclusively bind the TMD (Shimizu, Guo and Gardella, 2001), and binds in a manner that is strongly dependent on presence of a coupled G protein (Okazaki *et al*., 2008). Labeled M-PTH derivatives can therefore be used as tracers to measure the ability of ligands to engage the PTH1R in the G protein bound state (Dean *et al*., 2006).

Our cryo-EM data is consistent with the hypothesis that M-PTH predominantly binds the TMD. The C-terminal-most residue of M-PTH, Trp14, only extends to the top of TM1, and W14 π-stacks with F184^1.36^ in the same manner as does the analogous residue of LA-PTH (**Figure 3a**). The elongated sidechain of the homoarginine 11 residue, which enhances potency 30-fold compared to the native leucine of PTH_(1-14)_ (Shimizu *et al*., 2001), penetrates far into the TM1-TM2 cleft to contact Y245^2.72^ and to form electrostatic interactions with D241^2.68^. The two Aib residues, which collectively increase potency 100-fold (Shimizu, Guo and Gardella, 2001), form extensive contacts in the orthosteric pocket, including hydrophobic interactions with L368^5.44^ and M441^7.39^. Strikingly, we detected little to no density for the ECD (**Figure 1a**), indicating that it is very mobile, and this was also reflected in the 3DVA where density for the ECD was absent even at low contour (**Video S1**). This observation supports the hypothesis that class B1 GPCR ECDs are highly dynamic when they are not anchored by interaction of ligands, or other proteins such as receptor activity-modifying proteins (Josephs *et al*., 2021), that bridge the TMD and ECD (Yang *et al*., 2015; Cong *et al*., 2021).

### The PTH1R uses structural waters to bind ligands and recruit G protein

GPCRs employ rigidly bound water molecules to support tertiary structure, bind ligands, aid receptor activation, and recruit effector proteins (Venkatakrishnan *et al*., 2019). Within our sub-3Å resolution maps, we were able to place several waters in the atomic models, including some at sites that are topologically conserved with other class B1 GPCRs (Sun *et al*., 2020; Zhang *et al*., 2020). In the receptor’s intracellular cavity, a water molecule forms bridging hydrogen bonds between the sidechain of Y391^Gs^ and the conserved residue E302^3.50^. Y391^Gs^ also interacts with a water molecule via its backbone carbonyl (**Figure 3b; Table S1**). The sidechains of R219^2.46^ and H223^2.50^, two highly conserved residues, are linked by a water-mediated hydrogen bond. In each of these cases, analogous waters were observed in structures of the glucagon-like peptide-1 receptor (GLP-1R) bound to Gs (**Figure 3b**) (Zhang *et al*., 2020).

We observed three water molecules in the vicinity of the centrally located P^6.47^xxG^6.50^ motif. This critical motif facilitates the formation of a pronounced kink in TM6 that is stabilised by G protein recruitment (de Graaf *et al*., 2017). Notably, a water makes bridging hydrogen bonds to the backbones of G418^6.50^ and I371^5.47^ (**Figure 3c; Table S1**). Equivalent waters were observed in the case of the GLP-1R, as well as in the more distantly related calcitonin receptor (Cao *et al*., 2022), illustrating importance of waters in activation mechanisms, and stabilisation of the G protein coupled state that is conserved across class B1 GPCRs.

In the region where PTH interacts with the TMD, we were able to identify a number of sites for waters. Most of these sites are also apparent in the structures containing PTHrP or LA-PTH, but the higher resolution of the PTH containing map enabled further discrimination. Two waters, one hydrogen bonded with the backbone of Ser1 and one hydrogen bonded to the backbone of Glu4, mediate PTH’s interaction with F424^6.56^ and Y296^3.44^, respectively (**Figure 3d; Table S1**). In the orthosteric pocket, the interactions between Y245^2.72^ and K240^2.67^ are mediated by a water, and a further water was identified in the vicinity of ECL2, hydrogen bonded to Ser356^ECL2^ (**Table S1**). These waters likely contribute to affinity and specificity of ligands.

## Discussion

PTH1R-mediated signaling is a complex, multistep process (Ferrandon *et al*., 2009; Feinstein *et al*., 2011) that is not yet completely understood. In the context of therapeutic PTH1R agonists, the duration of signaling via Gαs and cAMP production is hypothesized to be a key parameter for pharmacodynamics (Cheloha *et al*., 2015; White *et al*., 2021). Duration of signaling is proposed, at least partially, to be related to peptide affinity for the receptor in the presence or absence of G protein (states referred to as R^0^ and R^G^, respectively) (Okazaki *et al*., 2008). The relatively poor ability of M-PTH to bind to the R^0^ state (Okazaki *et al*., 2008) can be rationalized by its lack of ECD engagement, an interaction which does not require G proteins (Pioszak and Xu, 2008). For the full-length peptides that do engage the ECD, an explanation for the marked differences in binding affinity for the R^0^ state (rank order LA-PTH > PTH > PTHrP > ABL) (Hattersley *et al*., 2016) was not immediately apparent from our cryo-EM data. This observation is unsurprising given that each of these peptides show comparable affinity in the R^G^ state (Hattersley *et al*., 2016), and we imaged the PTH1R in an R^G^ state. The peptide C-terminus can bind the ECD in the absence of G protein, however, the subtly altered configurations/dynamics of the ECD that we observed could manifest as differences in R^0^ affinity.

Full-length peptide agonists of the PTH1R bind in a two-step process (Castro *et al*., 2005), corresponding to sequential binding to the ECD and the TMD. Previous data suggested that plasticity of the peptide-TMD engagement is relevant for activity at the glucagon-like peptide-1 receptor (GLP-1R) (Cary *et al*., 2022; Deganutti *et al*., 2022), a related class B1 GPCR, where the N-terminus of some peptides become partially or fully unbound in 3DVA associated with outward movement of ECL3. This dynamic behaviour was commonly observed in peptides that have residues that can destabilise helical secondary structure of the activation domain of those peptides (Johnson *et al*., 2021; Zhang *et al*., 2021; Cary *et al*., 2022). In contrast, all PTH1R-peptide complexes in the current study exhibited stable interactions between the peptide N-terminus and the receptor core (**Video S1**), suggesting that this interaction might be generally more stable than for agonists of the GLP-1R. Alternatively, there may be greater allosteric coupling between the deep (fully engaged) binding state and G protein binding for the PTH1R, such that intermediate states where the G protein remains bound are not sufficiently stable to be captured; this could contribute to the sample heterogeneity observed for all PTH1R complexes under the conditions of the current study. Further work is required to address these hypotheses.

## Conclusion

The PTH1R is an important target for the treatment of osteoporosis and a vital mediator of serum calcium homeostasis. Here, we present insights into structural underpinnings for binding of diverse peptide agonists to the PTH1R. We show that active fragments of the two native hormones bind similarly to the TMD, consistent with measurements in assays of Gs activation. Our structure of the truncated agonist M-PTH demonstrates that the PTH1R extracellular domain is highly mobile in the absence of tethering by a bound peptide. This structural dataset, in particular our high resolution models where waters could be identified, complement recent advances to understand the structure and function of the class B1 receptor family (Cong *et al*., 2022).

While this work was being finalized, Zhang *et al*. and Kobayashi *et al*. made available reports detailing cryo-EM structures of PTH1R bound to PTH and ABL (Liu *et al*., 2022) and PTH and PTHrP (Kobayashi *et al*., 2022), respectively. Our structures of PTH, ABL, and PTHrP closely agree with those presented by the other groups, these three reports substantially bolster understanding for the molecular basis of PTH1R signaling.

## Lead contact

Further information and requests for resources and reagents should be directed to and will be fulfilled by the Lead Contact, Patrick Sexton.

## Materials availability

The unique reagent generated in this study (the HA-FLAG-3C-PTH1R-3C-His construct in pFastbac-Dual) is available from the lead contact.

## Data and code availability

The atomic coordinates and the cryo-EM density maps generated during this study are available at the protein databank (https://www.rcsb.org) and the electron microscopy databank (https://www.ebi.ac.uk/pdbe/emdb) under accession numbers PDB: XXXX PDB: XXXX, PDB: XXXX, PDB: XXXX, and PDB: XXXX, and EMDB entry ID EMD-XXXXX, ID EMD-XXXXX, ID EMD-XXXXX, ID EMD-XXXXX, and ID EMD-XXXXX, for the PTH, PTHrP, ABL, M-PTH and LA-PTH containing complexes, respectively.

This paper does not report original code. Any additional information required to reanalyze the data reported in this paper is available from the lead contact on request.

## SI Text

### PTH1R/Gs complex formation

Insect cell culture and protein expression were performed as previously reported (Liang *et al*., 2017). Briefly, cultures of *Trichoplusia ni* cells grown in ESF 921 insect cell culture medium were infected with baculoviruses encoding PTH1R, DNGαs (Liang *et al*., 2018), Gβ1 and Gγ2. After ∼48 h, the cultures were centrifuged (4000 *g*), and cell pellets from half-liter aliquots of culture were frozen and stored at -80°C until use.

*Trichoplusia ni* cell pellets (10-15 g) containing the desired proteins were thawed and suspended in buffer (40 mL, 30 mM HEPES, 50 mM NaCl, 2 mM MgCl_2_, and 5 mM CaCl_2_, pH 7.4). To this mixture were added 5-10 µM agonist peptide (or 100 µM in the case of M-PTH; as a stock solution in water), benzonase (2 µL) and cOmplete Protease Inhibitor Cocktail (1 or 2 tablets). After 30 min of stirring at room temperature, apyrase (5-10 µL) and nanobody35 (1 mg) were added to the mixture. The mixture was stirred for another 30 min at room temperature, and 10 mL of detergent solution (5% LMNG w/v and 0.3% w/v CHS in water) and 0.7 mL of NaCl (5 M) were slowly added. The resulting mixture was Dounce homogenized with a tight pestle. Buffer (40 mL, 30 mM HEPES, 50 mM NaCl, 2 mM MgCl_2_, and 5 mM CaCl_2_, pH 7.4) and NaCl (0.4 mL, 5M) were added, and the resulting mixture was stirred at 4°C for 1-2 h. Insoluble debris was removed by centrifugation (25,000 *g*, 20 min). The supernatant was filtered (glass fibre prefilter), and ∼3 mL anti-FLAG M1 affinity resin was added to the solution. The mixture was agitated at room temperature for 2 h and then the resin was loaded into a glass-fritted column. The resin was washed with 100 mL of wash buffer (20 mM HEPES, 100 mM NaCl, 2 mM MgCl_2_, 5 mM CaCl_2_, 2 µM agonist peptide (or 25 µM for M-PTH), 0.01% LMNG, and 0.00006% CHS), and protein was eluted with 10-20 mL of elution buffer (20 mM HEPES, 100 mM NaCl, 2 mM MgCl_2_, 0.2 mg/mL FLAG peptide, 10 mM EGTA, 2 µM agonist peptide (or 25 µM for M-PTH), 0.01% LMNG, and 0.00006% CHS). The eluent was concentrated to ∼500 µL using a 100 kDa MWCO centrifugal concentrator, filtered (0.22 µm), and injected onto a Biorad system equipped with a GE Superdex 200 Increase 1G/300 GL column, BioLogic dual flow, BioFrac fraction collector, and a Shimadzu RF10AXL fluorescence detector. The complex was resolved with a 0.5 mL/min flow of SEC buffer (20 mM HEPES, 100 mM NaCl, 2 mM MgCl_2_, 0.1 mM TCEP, 2 µM agonist peptide (or 25 µM for M-PTH), 0.01% LMNG, and 0.00006% CHS). Fractions (300 µL) were pooled and concentrated with either a 100 kDa or 50 kDa MWCO centrifugal concentrator for SEC peaks corresponding to PTH1R with or without G protein, respectively. Final samples were aliquoted, flash-frozen in liquid nitrogen, and stored at -80°C.

### SDS-PAGE

Samples were prepared for SDS-PAGE with a 1:1:1 mixture of sample, 10% (wt/vol) SDS (aq.) and Laemmli loading buffer containing 2-mercaptoethanol. Sample mixtures were not heated before loading onto gels. SDS–PAGE samples (10 µl) were loaded and run on Mini-PROTEAN TGX Precast 4–15% gels at 200V for 30 min. SDS–PAGE gels were stained with InstantBlue Coomassie stain (Abcam).

### Cryo-EM sample preparation

Freshly thawed sample of concentrated (3-4 mg/mL, 3 µL) PTH1R complex was applied to glow discharged (GloQube Plus, air chamber, 20 mA, 60 s, negative polarity) TEM grids (either 300 mesh UltrAuFoil r1.2/1.3 grids or 200 mesh gold-coated (Russo and Passmore, 2016) Quantifoil r1.2/1.3 grids). A Thermo Fisher Scientific MkIV Vitrobot operated at 4 °C and 100% humidity was used to blot (blot force 16-18, blot time 6-8s) and vitrify samples in liquid ethane. Specific parameters for sample preparation are available in Figure S2.

### Cryo-EM imaging

TEM grids with vitrified samples were clipped and loaded into either a Thermo Fisher Scientific Glacios (200 kV, Falcon 4) or a FEI Titan Krios (300 kV, Gatan K3 with Gatan energy filter) cryo-electron microscope. Automated data collection was performed using aberration free image shift (AFIS) as implemented in EPU 2 software (Thermo Fisher Scientific). Images were collected with a requested defocus range of ∼-0.6 to -1.5 µm. Specific parameters for each sample are available in Table S2.

For samples imaged on the Glacios microscope, the instrument was operated at a nominal magnification of 120,000 x in nanoprobe mode with a physical pixel size of 0.85 Å. An objective aperture (100 µm) and C2 aperture (50 µm) were used. Data was saved as electron-event representation (EER) files. The total dose applied was ∼50 e^-^/Å^2^ at a dose rate of ∼4.67 e^-^/px/s.

For samples imaged on the Titan Krios microscope, the instrument was operated at a nominal magnification of 105,000 x or 130,000 x in EFTEM nanoprobe mode with a physical pixel size of either 0.82 Å or 0.65 Å, respectively. The total dose applied was 60 e^-^/Å^2^ at a dose rate of either 8.821 or 12 e^-^/px/s. The energy filter was operated with a slit width of 10 eV, the Gatan K3 camera was operated in correlated double sampling (CDS) mode, and movies were saved as TIFF files with 60 subframes.

### Cryo-EM data processing

EER and TIFF files were pre-processed using the EPU_Group_AFIS.py script (https://github.com/DustinMorado/EPU_group_AFIS) for import into RELION 3.1.2. MotionCor2 as implemented in RELION 3.1.2 was used for patch motion correction (Zheng *et al*., 2017). Contrast transfer function (CTF) estimation was performed with either GCTF 1.18 (Zhang, 2016) or CTFFIND 4.1.14 (Rohou and Grigorieff, 2015) and micrographs were selected by estimated maximum resolution. Particle picking was performed with crYOLO 1.7.6 (Wagner *et al*., 2019). Multiple rounds of 2D classification and *ab initio* reconstruction were performed with cryoSPARC (Punjani *et al*., 2017). Particles were subjected to Bayesian polishing (Zivanov, Nakane and Scheres, 2019) as implemented in RELION 3.1.2. After additional rounds of 2D classification, non-uniform refinement with CTF corrections was applied using cryoSPARC. Local refinement using a mask excluding micelle and G protein density, resolution estimation, and 3D-varability analysis with 3 components were performed using the cryoSPARC software package (Punjani and Fleet, 2021). Specific processing information for each sample can be found in Figures S3-S4.

### Model building and refinement

By comparing consensus maps, we noted a small discrepancy in the calibrated pixel sizes (0.8817 Å/px or 0.878 Å/px for PTHrP and ABL or M-PTH containing complexes, respectively) for datasets collected on the Glacios TEM compared to those collected on the Krios TEM. For a better comparison among structures from the two instruments, we refined models for complexes collected on the Glacios TEM against maps rescaled to 0.85 Å/px. We deposited these rescaled maps, as well as the original maps, half-maps, and FSC curves, in the EMDB.

Models were built using the protein databank (PDB) entries 6NBF (Zhao *et al*., 2019) and 6×18 (Zhang *et al*., 2020) as starting points for the PTH1R and G protein heterotrimeric complex, respectively. Flexible fitting was performed using ISOLDE 1.3 (Croll, 2018) as implemented in UCSF ChimeraX 1.3 (Goddard *et al*., 2018) and manual refinement was performed using COOT 0.9.6 (Emsley *et al*., 2010). Real space refinement and validation was performed Phenix 1.19.2 (Adams *et al*., 2010). All figures containing maps or models were generated using UCSF ChimeraX 1.3.

### cAMP accumulation assays

COS-1 cells were transiently transfected, using 1mg/ml polyethylenimine (PEI) Max (m.w. 40,000 Polysciences, Warrington, PA, USA). Cells were transfected with 100 ng/well of receptor. DNA and PEI, each diluted in 10µl of 150mM NaCl per well, were combined in a 1:6 ratio before being briefly vortexed and incubated for 15 min at room temperature. The transfection mixture was added dropwise to the cells and subsequently seeded into plates at 30,000 cells/well into 96-well clear plates (Corning) and incubated at 37°C in 5% CO_2_.

After 48 h cells were washed and changed into stimulation buffer (phenol-red free DMEM containing 0.1% w/v BSA and 0.5 mM IBMX (3-isobutyl-1-methylxanine), pH 7.4) and incubated for 30 min in 37°C, 5% CO_2_ before cells were stimulated with peptide. After the 30 min the reaction was terminated by the aspiration of the buffer and addition of 50 µl of ice-cold absolute ethanol. Upon the evaporation of ethanol, the cells were lysed with 75 µl of lysis buffer (5 mM HEPES, 0.1% w/v BSA, 0.3% Tween20, pH 7.4). The concentration of cAMP in the lysates was detected with the LANCE TR FRET kit (Perkin Elmer, Waltham, MA, USA). The plates were read on an Envision multi-label plate reader (Perkin Elmer) and values converted to an absolute concentration of cAMP using a cAMP standard curve detected in parallel.

### cAMP time course assay

HEK293A WT cells stably expressing an Epac-cAMP sensor (CAMYEL (Jiang *et al*., 2007)) were transiently transfected with FLAG-tagged PTH1R receptor using a standard PEI-based protocol at a ratio of 1 µg DNA: 6 µl PEI. Cells were seeded post-transfection at a density of 25,000 cells per well into a white, poly-D-lysine coated, 96-well plate and incubated for 48 h at 37ºC in 5% CO2. Growth media (DMEM + 5% FBS) was replaced with imaging buffer (phenol-free DMEM containing 1% (w/v) ovalbumin) and incubated at 37ºC in 5% CO2 for 1 h. Substrate (coelentazine-h) was added to a final concentration of 10 µM and incubated for 10 min before the start of the assay. BRET1 measurements were recorded using a PHERAstar plate reader (BMG labtech) at a rate of 1 cycle/114s at 37ºC. Ligands were added, at increasing concentrations diluted in the imaging buffer, at cycle 5, in duplicate. Data were subjected to baseline subtraction of the mean BRET ratio before ligand addition, then baseline subtraction to vehicle treated wells, and finally normalized to a 10 µM forskolin positive control.

### TRUpath assay

HEK293a cells were transiently transfected with FLAG-tagged PTH1R receptor and TRUpath components (αS-L-Rluc8, β1 and γ1-GFP) using a standard PEI-based protocol at a ratio of 1 µg DNA: 6 µl PEI. TRUpath components (Olsen *et al*., 2020) were added at a ratio of 1:1.5 µg receptor DNA. Cells were seeded post-transfection at a density of 25,000 cells per well into a white, poly-D-lysine coated, 96-well plate and incubated for 48 h at 37ºC in 5% CO_2_. Growth media (DMEM+ 5% FBS) was replaced with imaging buffer (10mM HEPES, 1x HBSS + 1% (w/v ovalbumin) and incubated at 37ºC in 5% CO_2_ for 1 h. Substrate (Prolume Purple) was added to a final concentration of 1 µM and incubated for 10 min before the start of the assay. BRET2 measurements were recorded using a PHERAstar plate reader (BMG labtech) at a rate of 1 cycle/114s at 37ºC. Ligands were added at increasing concentrations, diluted in the imaging buffer at cycle 5, in duplicate. Data was normalised to the mean baseline BRET ratio and then vehicle treated wells.

## Author Contributions

B.P.C., P.M.S, and D.W. contributed to conceptualization. B.P.C. expressed protein and purified complexes. Y.J. and I.C.R. prepared baculovirus. E.J.G. and M.M.F. performed assays. B.P.C. and M.J.B. prepared cryo-EM samples and acquired EM data. B.P.C., M.J.B., and S.J.P. processed single-particle cryo-EM data. B.P.C built atomic models. B.P.C., E.J.G, and S.J.P. prepared figures. All authors reviewed and/or revised the paper.

## Acknowledgements

Cryo-EM samples were imaged at the at the Bio21 Advanced Microscopy Facility (The University of Melbourne) and at the Monash Ramaciotti Centre for Cryo-Electron Microscopy. High-performance computing was supported by Monash MASSIVE. P.M.S. is a Senior Principal Research Fellow (#1154434) and D.W, a Senior Research Fellow (#1155302) of the National Health and Medical Research Council of Australia. P.M.S is the Director and D.W. the Monash University Node leader of the Australian Research Council Industrial Transformation Training Centre for Cryo-electron Microscopy of Membrane Proteins (#IC200100052). This work was funded in part by a NHMRC Program Grant (#1150083) to P.M.S.

## Supplemental Figure Legends

**Figure S1:**
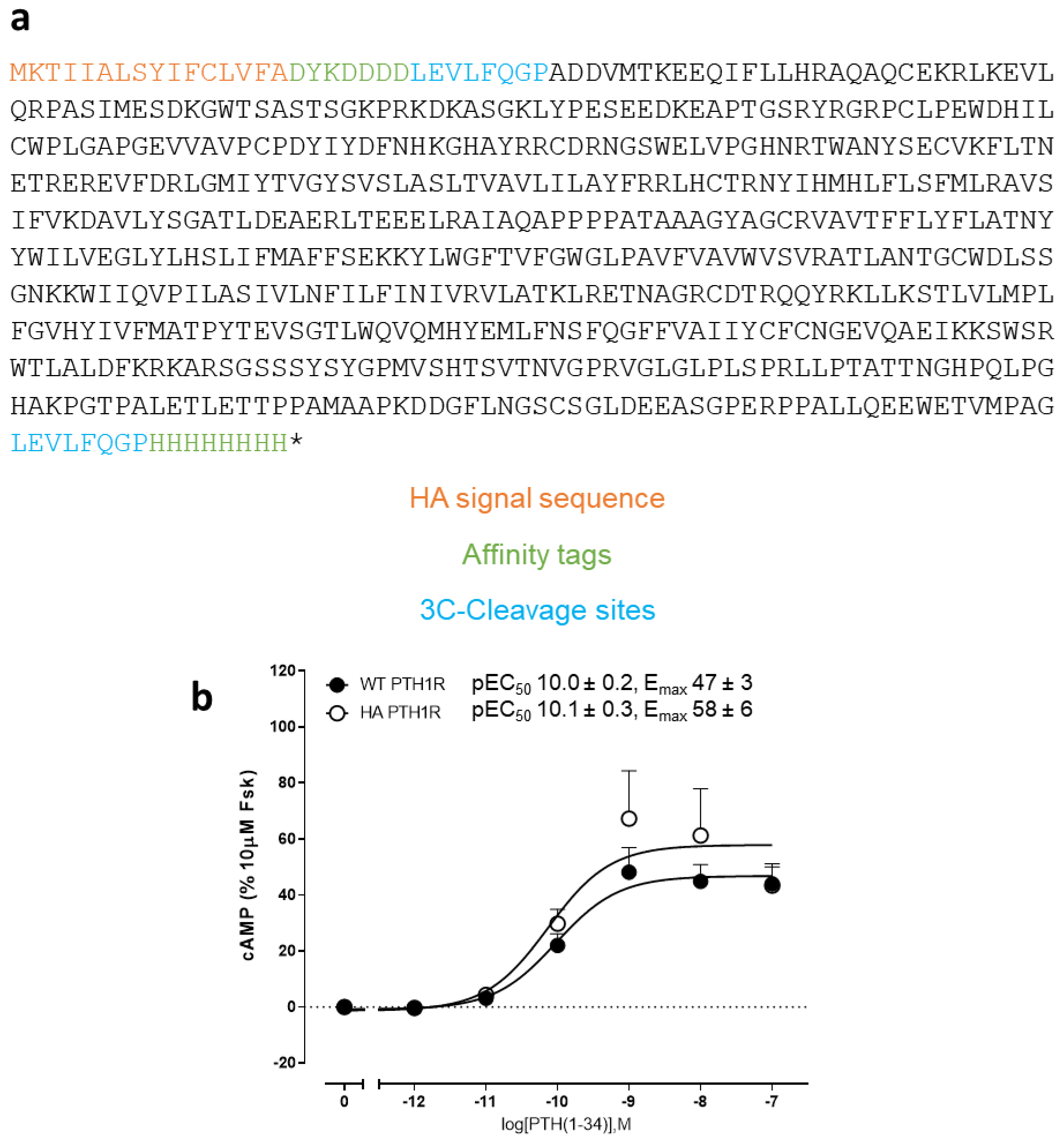
(**a**) Amino acid sequence of the PTH1R construct used for determination of cryo-EM structures. Orange text indicates the HA signal peptide, green text indicates affinity tags, and blue indicates 3C-cleavage sites. (**b**) Concentration response curve of wild-type PTH1R compared to the modified receptor used for structure determination in production of cAMP in response to PTH(1-34). 100% represents cAMP production in the presence of 10 µM forskolin (Fsk). Errors indicate standard error of the mean. N = 3.

**Figure S2:**
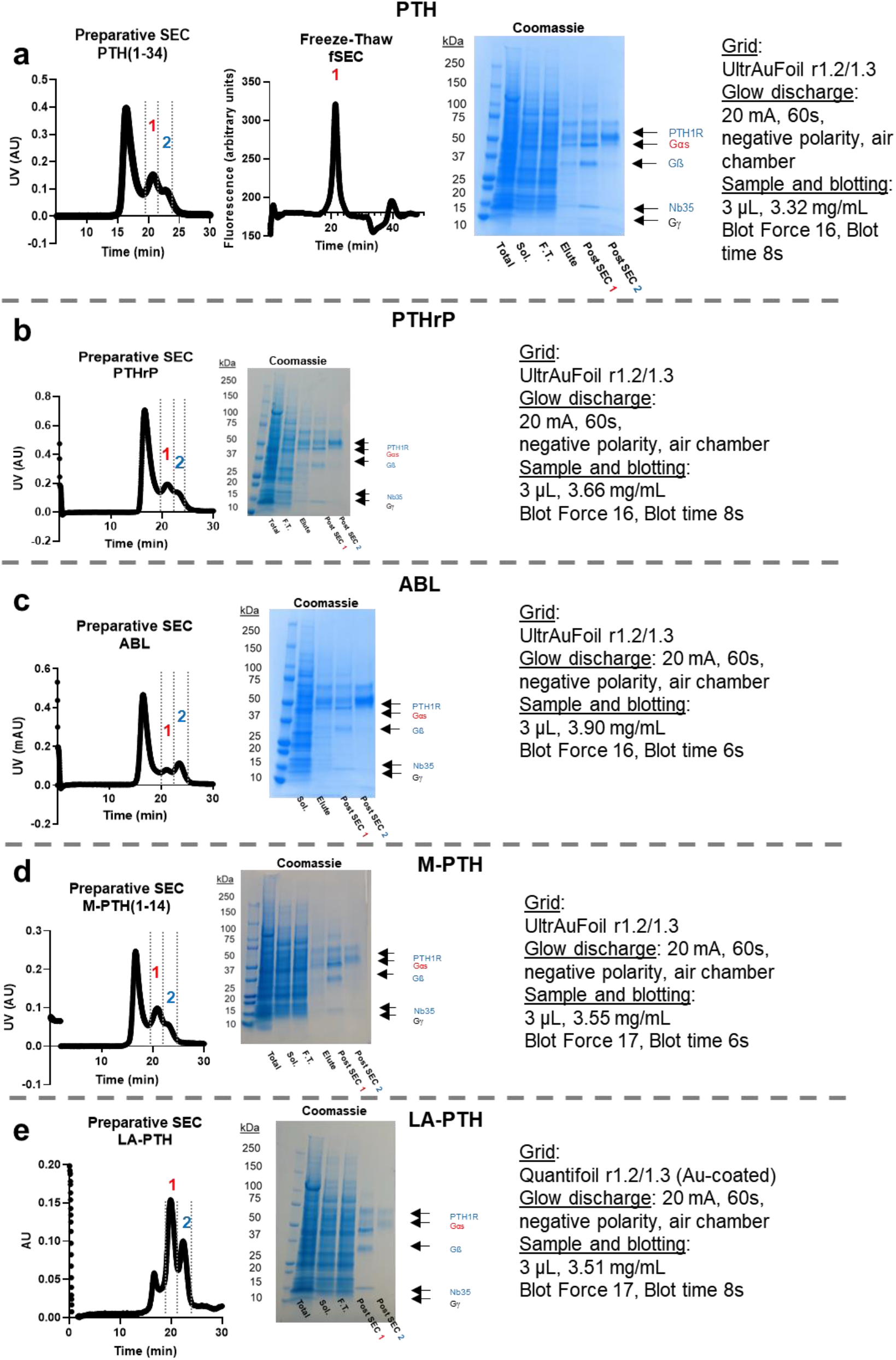
Sample preparation and analysis for PTH1R complexes containing: PTH (**a**), PTHrP (**b**), ABL (**c**), M-PTH (**d**), and LA-PTH (**e**). **Left**: SEC chromatographs. **Middle**: SDS-PAGE gels with Coomassie staining; **Right**: cryo-EM grid preparation conditions. Sol. Indicates solubilized fraction, F.T. indicates anti-FLAG column flow through. Post SEC 1 and Post SEC 2 indicate samples of pooled fractions taken from the preparative SEC with approximate retention times as labeled in the chromatographs. In each case, cryo-EM samples were prepared from the sample labeled Post SEC 1. For the sample containing PTH, the sample was flash-frozen, thawed, and reinjected onto the SEC for analysis (labeled as Freeze-Thaw fSEC in panel **a**).

**Figure S3:**
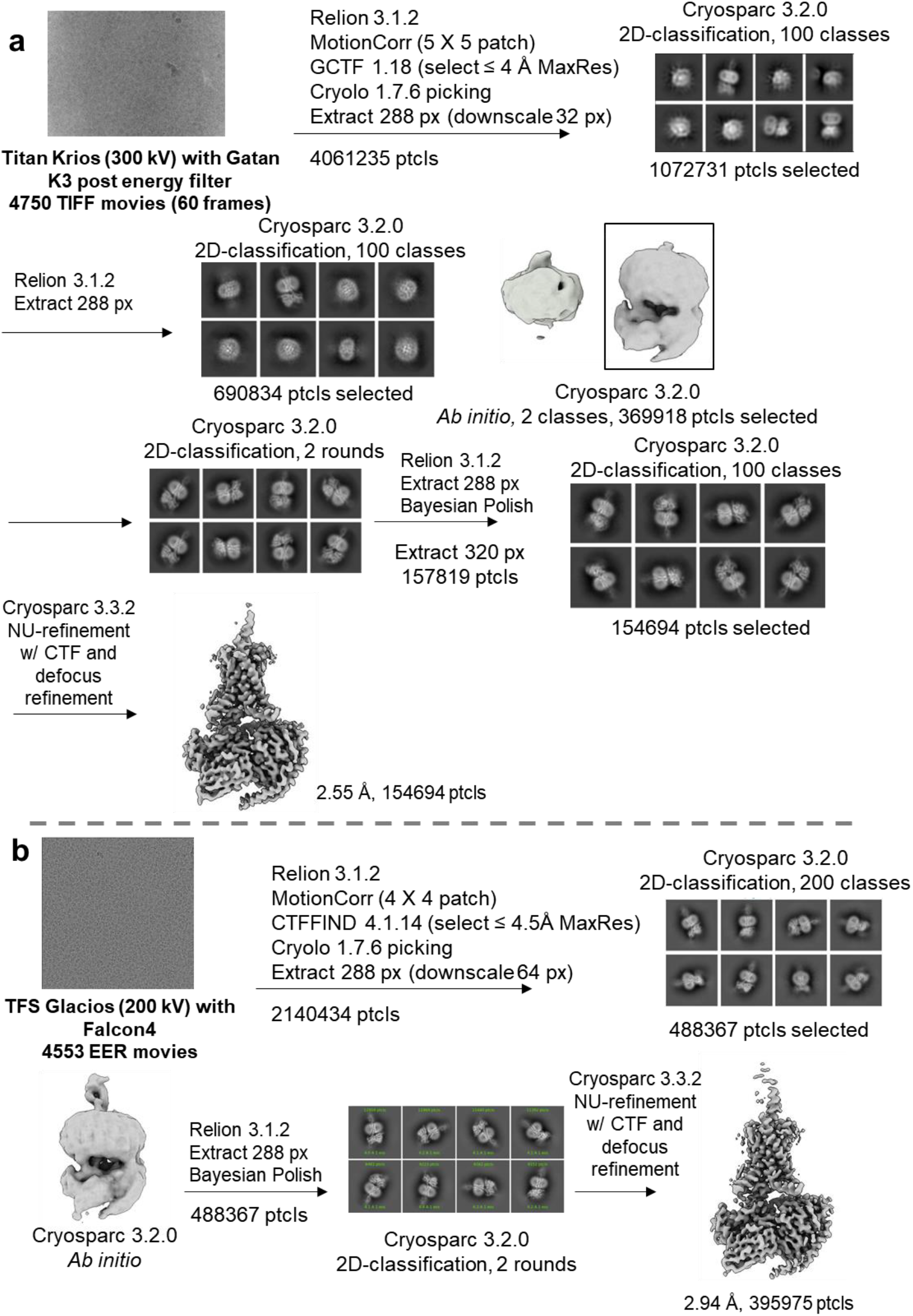
Overview of the processing pipeline for cryo-EM data of PTH1R complexes with PTH (**a**) and PTHrP (**b**).

**Figure S4:**
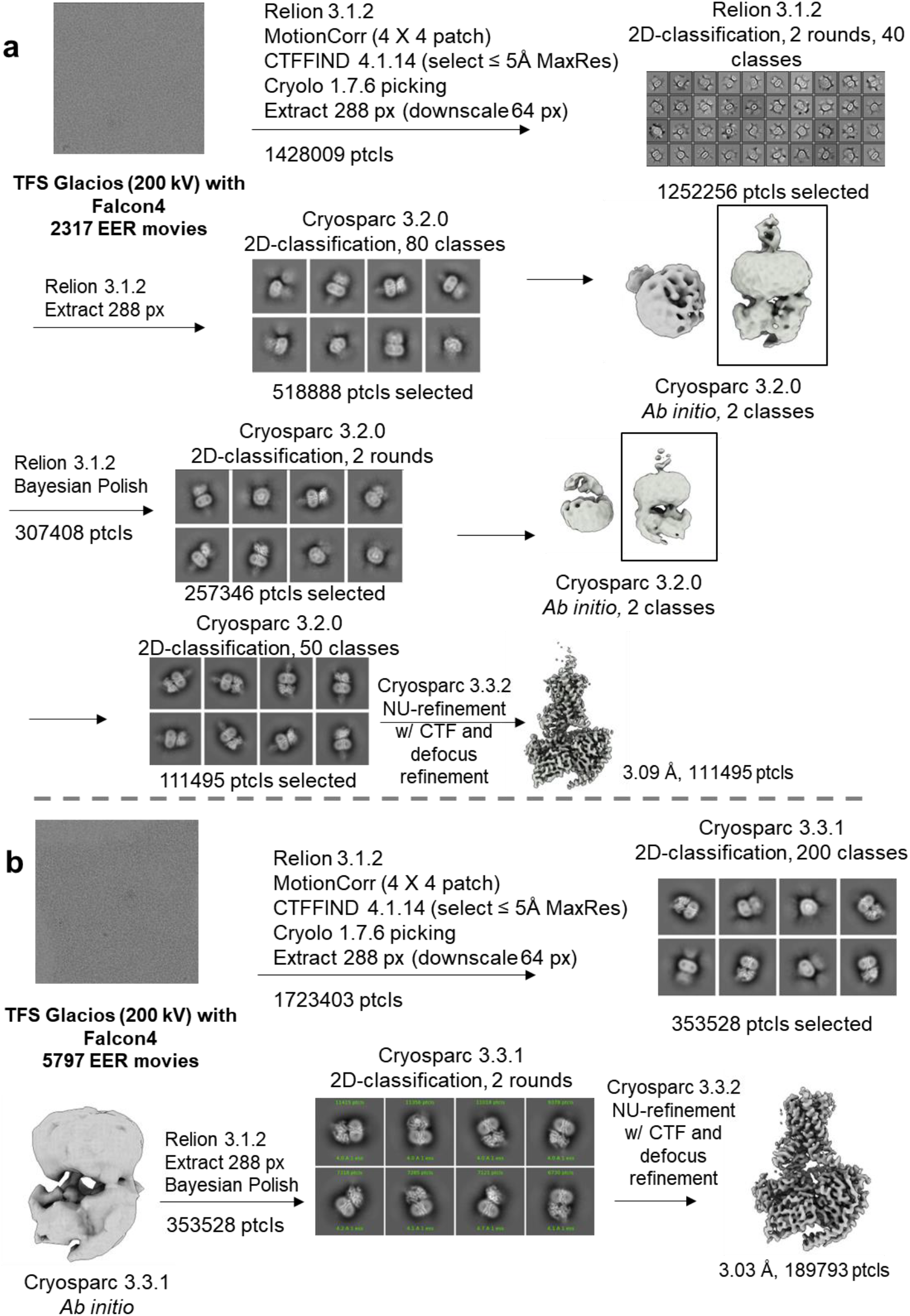
Overview of the processing pipeline for cryo-EM data of PTH1R complexes with ABL (**a**) and M-PTH (**b**).

**Figure S5:**
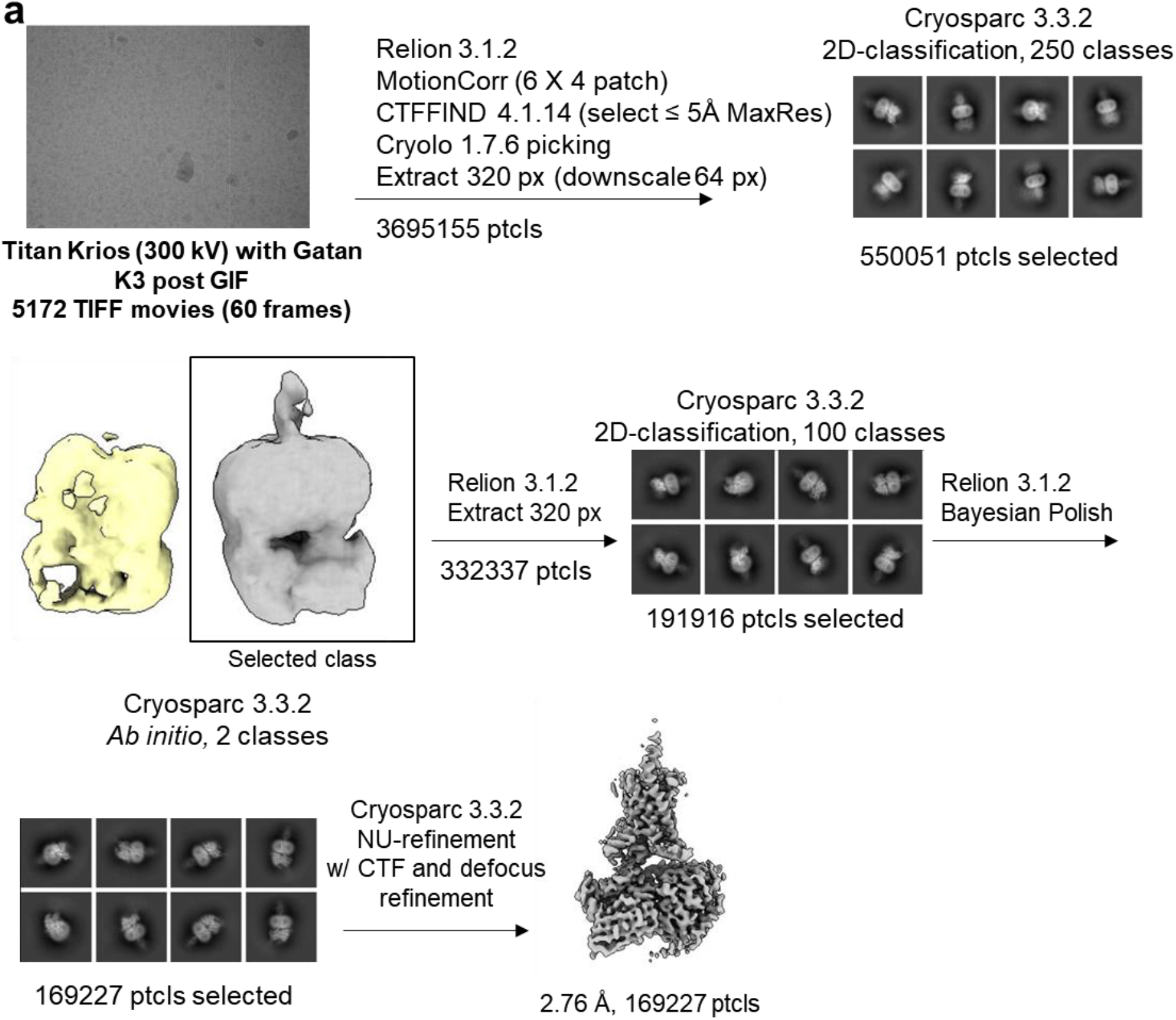
Overview of the processing pipeline for cryo-EM data of PTH1R complex with LA-PTH.

**Figure S6:**
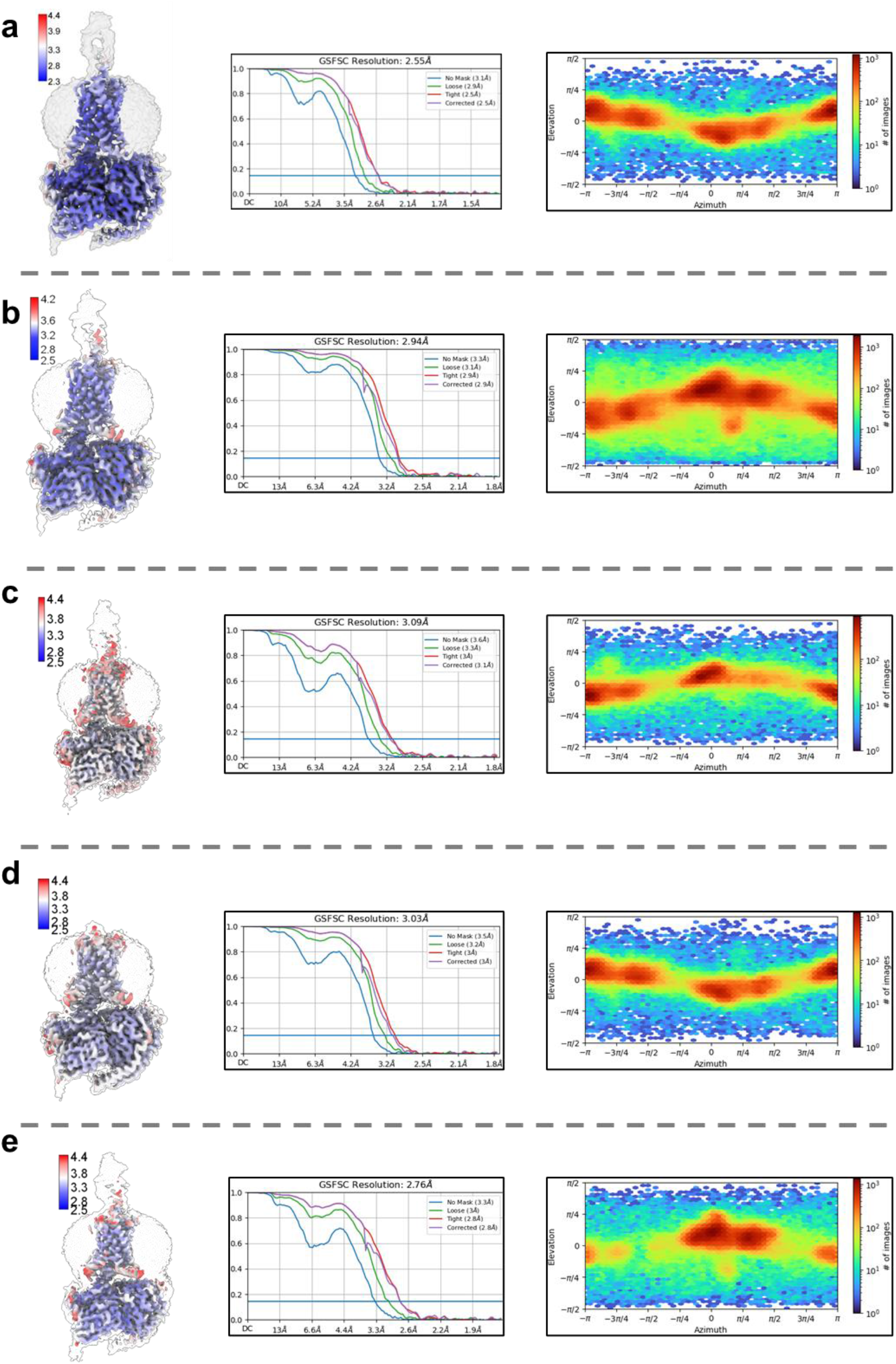
Local resolution estimates (**left**), Fourier shell correlation (**middle**), and particle viewing direction distribution plots (**right**) for samples containing PTH (**a**), PTHrP (**b**), ABL (**c**), M-PTH (**d**) and LA-PTH (**e**).

**Figure S7:**
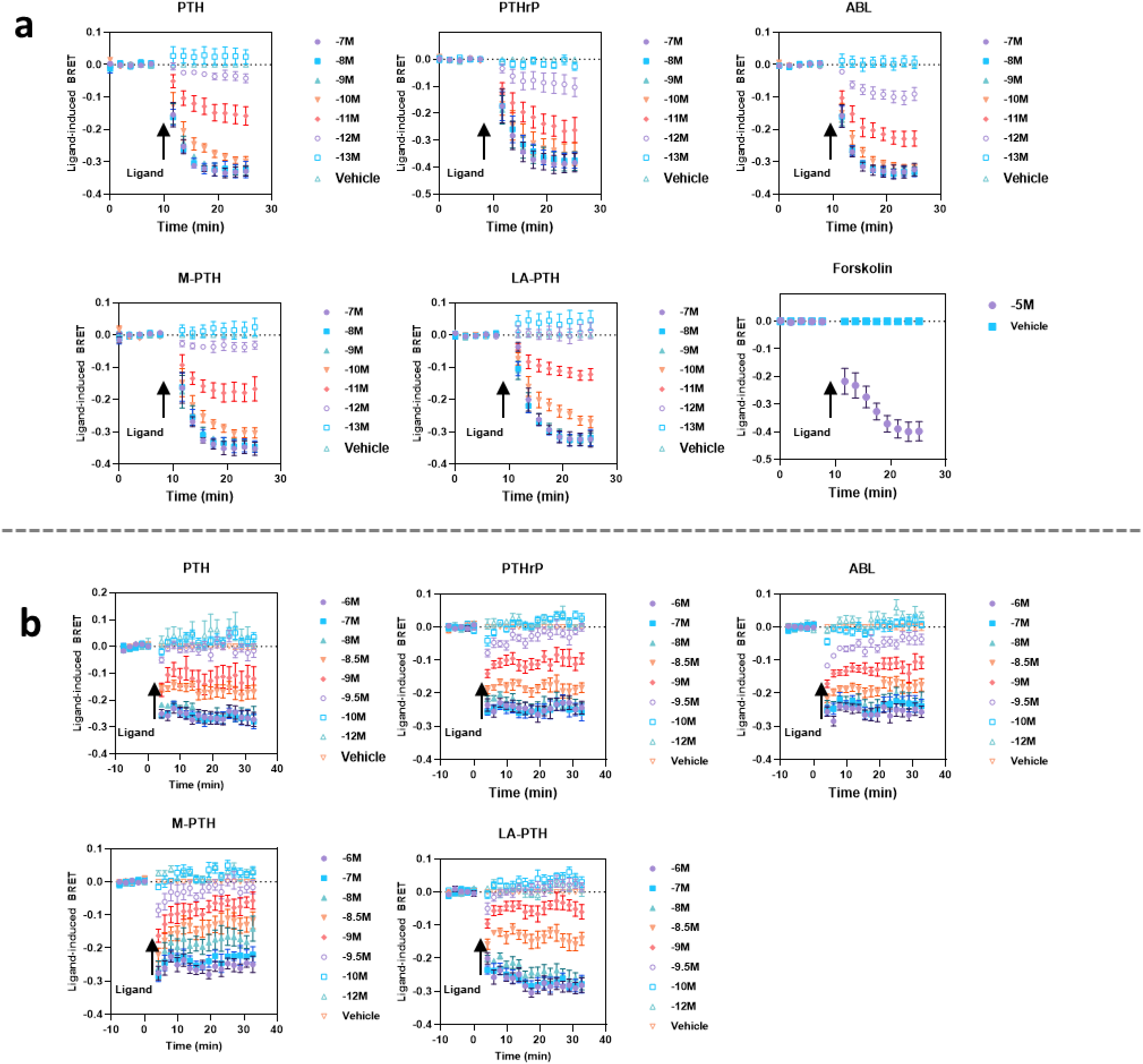
Kinetics of ligand induced PTH1R activation. (**a**) cAMP production as measured by the CAMYEL BRET biosensor in HEK293 cells transiently expressing the PTH1R. N = 3. (**b**) Gs (long variant) activation as measured by the TruPath BRET biosensor in HEK293 cells transiently expressing the PTH1R and sensor components. N = 4. Ligand was added at the timepoints indicated by arrows. Experimental uncertainties represent the standard error of the mean.

**Figure S8:**
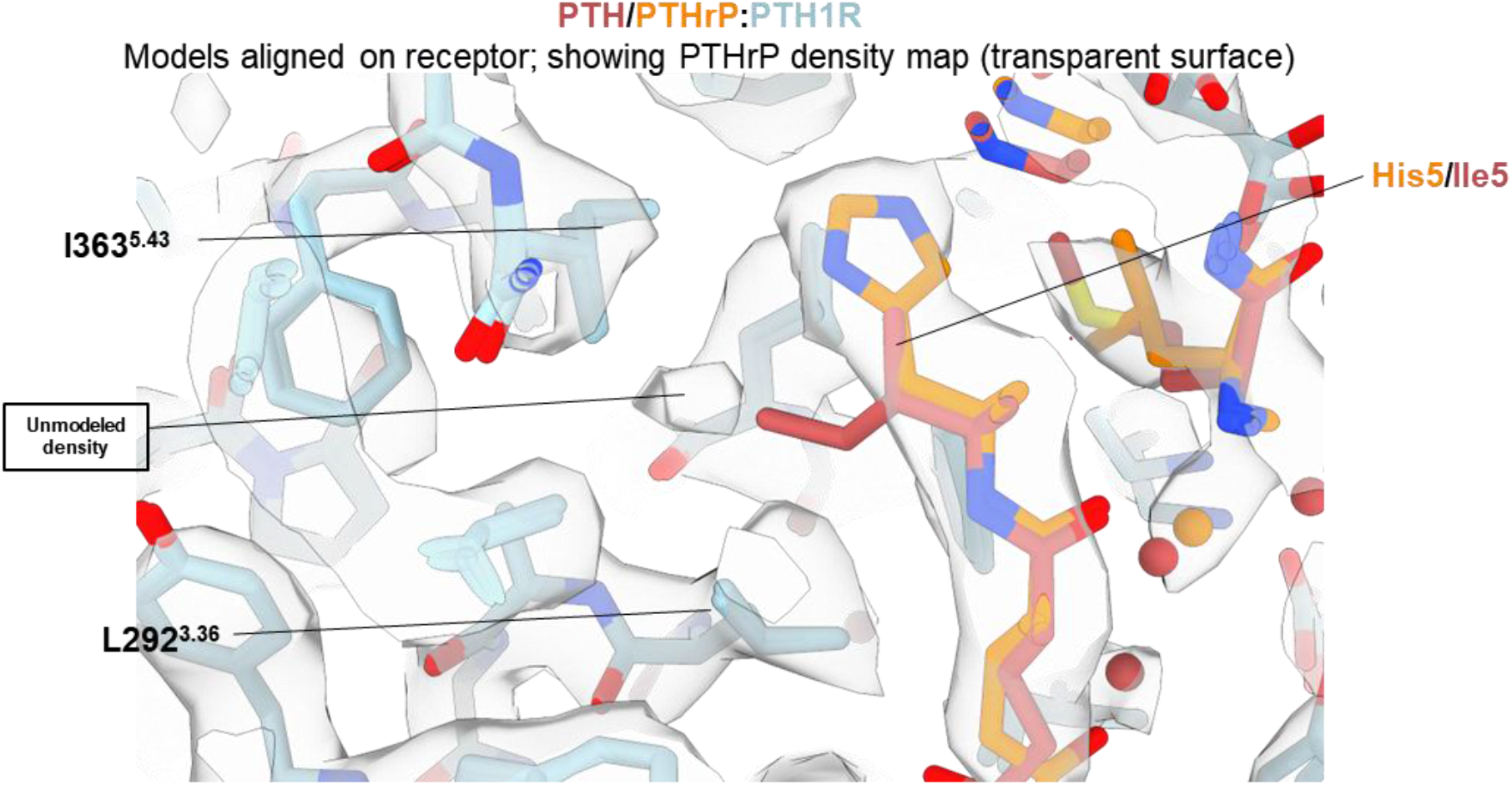
A comparison of the location of peptide residue 5 of PTH and PTHrP, shown in orange and red stick representation, respectively, in the PTH1R binding site. The cryo-EM density map for the PTHrP-containing map is shown as a light grey transparent surface.

## Supplementary Tables

**Table S1:**
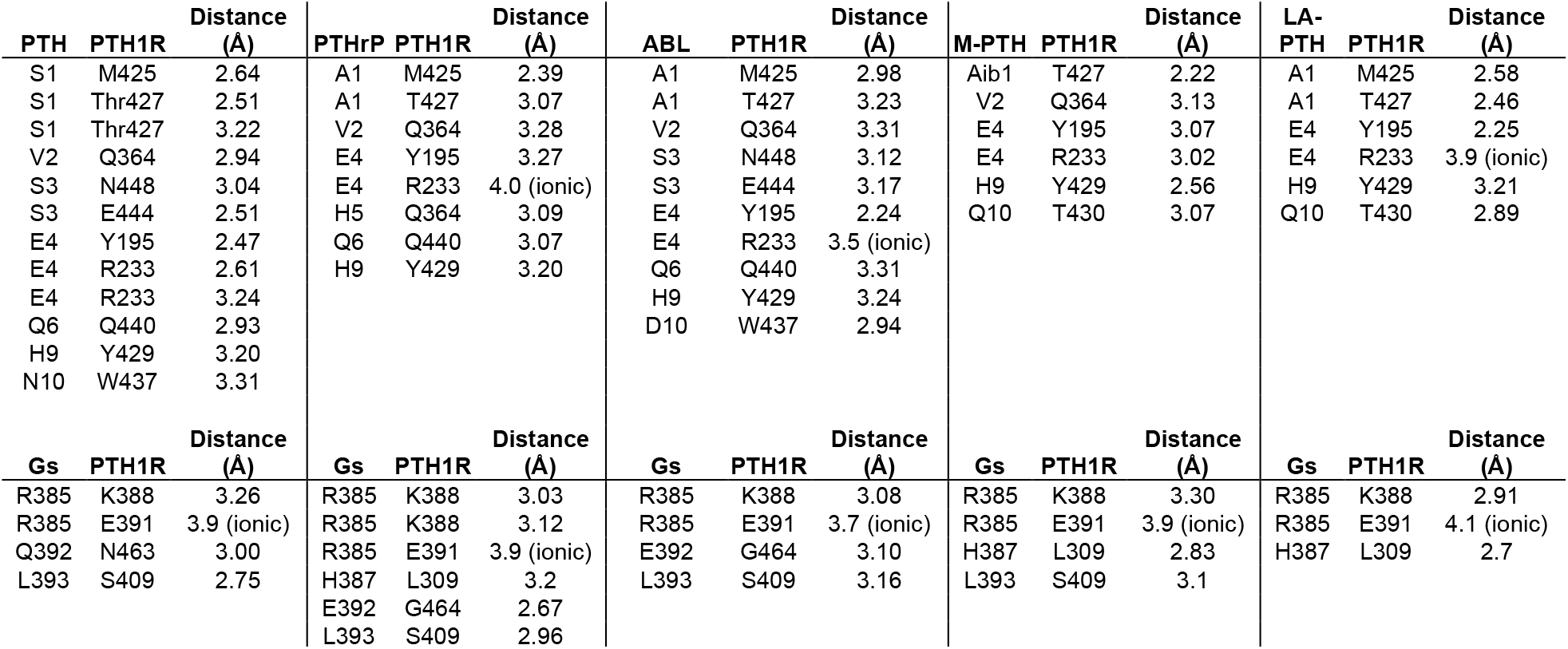
N-terminal peptide/receptor and C-terminal G protein/receptor polar interactions for each complex. Hydrogen bonds and ionic interactions between PTH1R and peptide agonist N-terminal regions as determined by LigPlot+ v2.2.5 (Laskowski and Swindells, 2011). Distances for ionic interactions were determined using PyMol 2.4.1.

**Table S2:**
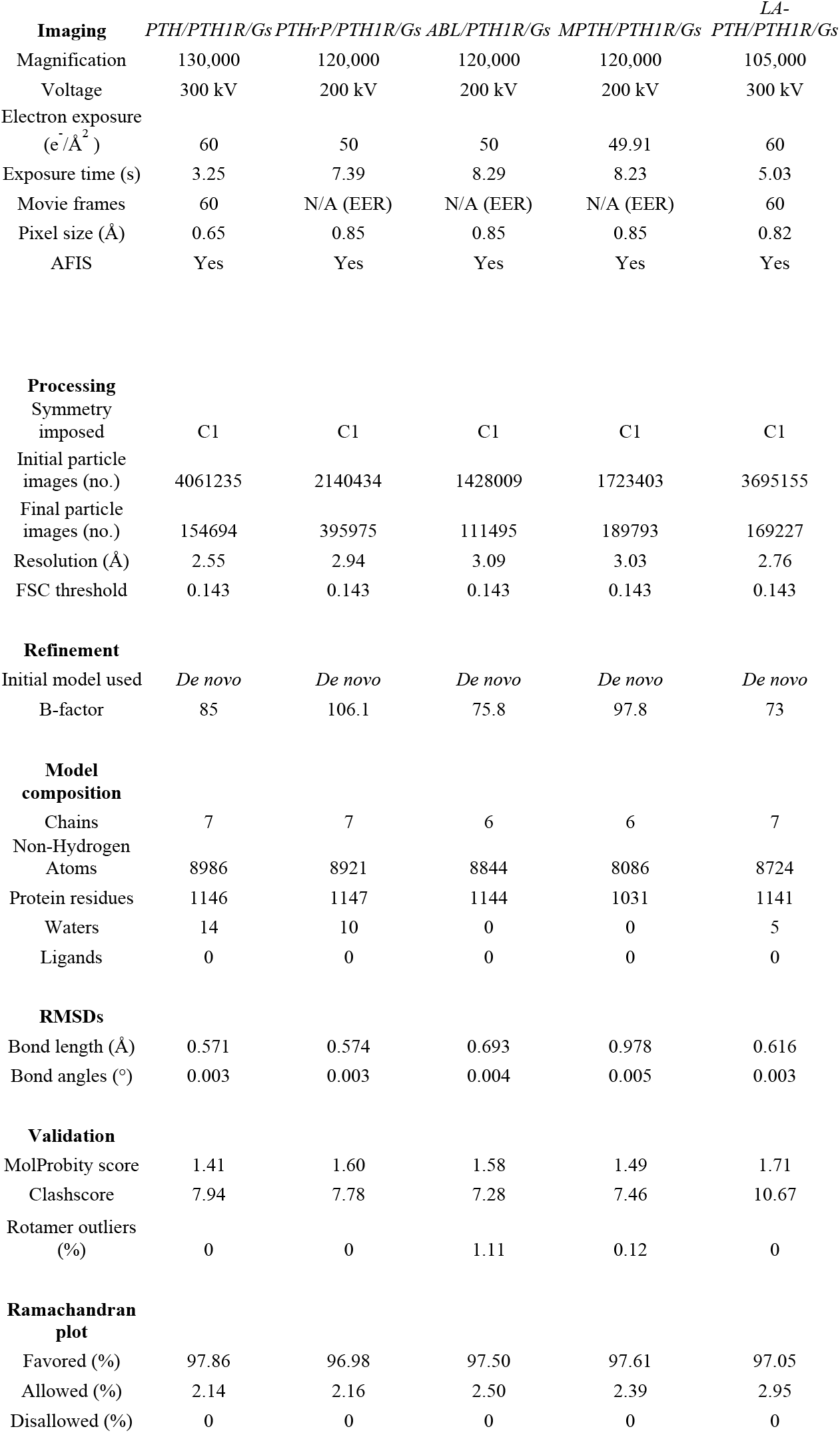
Cryo-EM imaging, refinement, and model parameters.

**Table S3:** Materials

| <u>Item/Reagent</u> | <u>Vendor</u> | <u>Item #</u> | <u>Notes</u> |
| --- | --- | --- | --- |
| ESF 921™ Insect Cell Culture Medium | Expression Systems | 96-001-01 |  |
| Peptide synthesis (PTH, PTHrP, M-PTH, Abaloparatide) | GL Biochem (Shanghai) Ltd | - |  |
| Peptide synthesis (LA-PTH) | Genscript | - |  |
| 4–15% Mini-PROTEAN® TGX™ Precast Protein Gels | BioRad | 4561086 |  |
| Anti-FLAG M1 Affinity Resin | Millipore Sigma | A4596 |  |
| EGTA | Combi-Blocks | QB-6401 |  |
| Precision Plus Protein™ Dual Color Standards | BioRad | 1610374 |  |
| Apyrase | New England Biolabs | M0398L |  |
| Lauryl Maltose Neopentyl Glycol (LMNG) | Anatrace | NG310 |  |
| Cholesteryl Hemisuccinate Tris Salt (CHS) | Anatrace | CH210 |  |
| Benzonase® Nuclease HC, Purity > 90% | Millipore Sigma | 71205 | 250 U/μl |
| cOmplete™, Mini Protease Inhibitor Cocktail | Roche (Millipore Sigma) | 11836153001 |  |
| Amicon Ultra Centrifugal Filter (MWCO, 100 kDa) | Millipore Sigma | UFC910024 |  |
| UltrAuFoil R1.2/1.3, 300 mesh grids | Electron Microscopy Sciences | Q350AR13A |  |
| Quantifoil R1.2/1.3, 200 mesh, Cu grids | Electron Microscopy Sciences | Q2100CR1.3 |  |
| DMEM | Thermo Fisher | 11995065 |  |
| HBSS | Thermo Fisher | 14065056 |  |
| Coelentazine-h | Nanolight Tech | 3031 |  |
| Prolume purple | Nanolight Tech | 369 |  |
| Forskolin | Millipore Sigma | F6886 |  |

## Supplementary Video Legend

**Video S1:** A compilation of movies showing the results of 3D-variability analysis. The series of 20 density maps for each complexes containing PTH, ABL, LA-PTH, M-PTH and PTHrP are shown oscillating, first at high threshold and then at lower threshold. The three variability components (labeled 0-2) for each complex are shown in gray, pink, and blue, respectively.

